# Elastogenesis by adventitial progenitors acquiring a smooth muscle cell phenotype following aortic dissection

**DOI:** 10.64898/2026.06.11.731783

**Authors:** Sohei Ito, Preet Patel, Taishi Inoue, Ruizhi Wang, Yuriko Katsumata, Hong S. Lu, Kenji Okada, Alan Daugherty, Hisashi Sawada

## Abstract

Following aortic dissection (AD), there is a sustained risk of vascular complications, progressive false lumen aneurysm formation, and rupture. However, no effective therapy exists to prevent these complications, highlighting the need to elucidate the pathophysiology following AD. Elastic fibers are crucial for maintaining aortic wall integrity but are thought to have limited regenerative capacity once disrupted during AD. This study defined that elastic fibers were newly generated in the false lumen wall following AD in humans and mice. In human ADs, new elastic fibers were observed in the false lumen wall 6 months after onset. In mice with descending AD induced by β-aminopropionitrile (BAPN), elastin mRNA was markedly upregulated in the chronic phase following AD, accompanied by elastic fiber formation. These fibers coincided with smooth muscle cell (SMC) markers within the false lumen wall. Of note, lineage tracing studies demonstrated that these cells were not derived from resident SMCs but adventitial progenitor cells. In vitro experiments further demonstrated that adventitial progenitor cells produced elastic fibers while expressing SMC markers. Collectively, these findings suggest that adventitial progenitor cells differentiate into elastogenic SMC-like cells, contributing to false lumen remodeling through de novo elastic fiber formation following AD.

## INTRODUCTION

Aortic dissection (AD) is a life-threatening vascular disease characterized by an intimomedial tear, resulting in blood entering the aortic media and forming a structurally fragile false lumen at a high risk for rupture and sudden death (1–3). AD is clinically classified into two types based on its location according to the Stanford classification: Type A AD involves the ascending aorta and type B AD is confined to the descending aorta. In the acute phase, treatment is selected based on disease type. Type A AD requires emergent surgical intervention, whereas type B AD is generally managed by pharmacological control of heart rate and blood pressure, unless patients progress into rapid dilatation, impending rupture, or organ malperfusion. Although the early mortality of type B AD is relatively lower, the false lumen can progressively undergo structural degeneration over time, leading to aneurysmal dilatation and rupture. Consequently, up to 60% of Type B AD patients eventually require surgical intervention, such as aortic replacement or stent graft implantation, due to these vascular complications (4). Despite its clinical significance, vascular complications following AD caused by false lumen degeneration have received limited attention in preclinical research. Most existing studies have focused on mechanisms of disease initiation, rather than its sequela (5, 6). Importantly, it is clinically challenging to predict the onset of AD in most cases. Thus, there is a critical need to elucidate the mechanisms related to false lumen degeneration to prevent post-AD complications.

Elastic fibers confer the aorta with compliance and an ability to store strain energy elastically, thus enhancing the conduit function under arterial pulsation (7–10). Synthesis of elastic fibers occurs in mid-gestation through the early postnatal phase of life (11–15). A previous study using radiocarbon labeling reported that the half-life of elastic fibers in humans is estimated to be 40-174 years (Mean: 74 years) (16). Therefore, elastic fibers are believed to be stable over the life span and not to regenerate in adulthood under normal conditions (8, 17, 18). While increased elastin mRNA abundance has been reported in aortic aneurysms of several mouse models (19–21), no study has demonstrated actual regeneration of elastic fibers in adult aortas. Thus, identifying mechanisms governing elastic fiber re-synthesis in the mature aorta provides a conceptual breakthrough in the extracellular matrix biology, with broad implications for the treatment of degenerative aortic diseases.

Fragmentation of elastic fibers is a prominent feature of false lumen degeneration in AD (22–24). As elastic fibers are essential for aortic integrity, their resynthesis is potentially beneficial in disease conditions to compensate for structural impairment. Therefore, we hypothesized that elastic fibers are newly generated following AD and mitigate post-AD complications. In the present study, we investigated whether elastic fibers are synthesized de novo in the dissected aortic wall after AD onset and explored the underlying mechanisms. We found that elastic fibers are newly formed within the false lumen wall during the chronic phase of AD in both humans and mice. Lineage tracing and spatial transcriptomic analyses combined with in vitro experiments uncovered the contribution of adventitial progenitor cells to de novo elastic fiber formation, with the differentiation into smooth muscle cell (SMC)-like cells in the dissected aortic wall.

## METHODS

### Sex as a Biological Variable

Human aortic samples were obtained exclusively from male patients because only male specimens were available from the surgical cohort, rather than due to intentional exclusion of female patients. Therefore, whether these findings are generalizable to females remains to be determined. Mouse experiments primarily used male mice in BAPN-induced AD studies, as previous reports demonstrated no significant sex differences in the incidence of AD or rupture in this model (25). For lineage-tracing experiments requiring in-house breeding of specific transgenic mouse lines, both male and female mice were included to reduce unnecessary animal sacrifice.

### Human Aortic Samples

Descending aortas were obtained from patients undergoing surgical repair of false lumen aneurysms at Kobe University. Aortic tissues were fixed with formalin (10% wt/vol) and stored in ethanol (70% vol/vol). Subsequently, tissues were embedded in paraffin blocks and sliced into 5 μm sections.

### Mice

C57BL/6J (#000664), *Rosa26*-mTmG (#007676), *Myh11*-CreER^T2^ (#0037658), and *Gli1*-CreER^T2^ (#007913) mice were purchased from The Jackson Laboratory. *Acta2-*CreER^T2^ and *Rosa26-*Rainbow mice were kindly provided by Dr. David A. Dichek (University of Washington) and Dr. Nicholas J. Leeper (Stanford University), respectively. Mice were housed in individually ventilated cages (5 mice/cage) on a light:dark cycle of 14:10 hours. Cage bedding consisted of Teklad Sani-Chip (#7090A, Envigo). Animals were fed a standard rodent diet (#2918, Envigo) and given either regular drinking water or water supplemented with β-aminopropionitrile (BAPN) (0.5% wt/vol, #A0796-500G, TCI Co.), both available ad libitum. BAPN was started when mice were 4 weeks of age. Fresh drinking water with BAPN was replaced twice each week. Necropsy was performed on any mice that died before the study endpoint to determine the cause of death.

### Histological Analyses

For lineage tracing, the descending aortas were fixed in paraformaldehyde (4% wt/vol) for 12 hours, followed by immersion in sucrose (30% wt/vol) for 24 hours. The tissues were then embedded in optimal cutting temperature compound (#4583, Sakura Finetek) and sectioned at a thickness of 10 μm. The sections were washed three times with phosphate-buffered saline and mounted using an anti-fade mounting medium containing DAPI (#ab104139, abcam). Fluorescent images were captured at 20x magnification using the tile image function on a Nikon AXR inverted confocal microscope. For other histological assays, descending aortas were immersed in paraformaldehyde (4% wt/vol), followed by incubation in ethanol (70% vol/vol), and subsequently embedded in paraffin. Paraffin-embedded sections (5 µm) were deparaffinized using limonene (#183164, Millipore-Sigma). Hematoxylin and eosin (#26043-06, Electron Microscopy Sciences; #ab246824, abcam) and Verhoeff iron hematoxylin staining (#041955.22, Thermo Fisher Scientific; #0512, VWR; #3803815, Leica Biosystems; #3120-32, Bicca) were performed as described previously (26). Immunostaining for Sca1 and ER-TR7 was performed on fixed frozen sections, whereas the other immunostaining was conducted on paraffin-embedded sections. Immunostaining was performed using primary and secondary antibodies listed in the Major Resource Table in the Supplemental Material. NovaRed (#SK-4805, Vector) was used as a chromogen. Histological images were captured using the Nikon AXR inverted confocal microscope or an Axio Scan Z7 (Zeiss).

### In Situ Hybridization

The distribution of mouse and human elastin mRNA in dissected aortas was examined by RNAscope® following the manufacturer’s instructions (Advanced Cell Diagnostics). After fixation using paraformaldehyde (4% wt/vol) or formaldehyde (10% wt/vol), aortic tissues were processed, embedded in paraffin, and cut at a thickness of 5 µm. Subsequently, sections were deparaffinized using xylene, followed by 2 washes with absolute ethanol. Target retrieval (#322000) was performed for 15 min at 100°C, followed by a protease (#322331) incubation step for 15 min at 40°C. Target mRNA was hybridized with mouse *Eln* (#319361) or human *ELN* probe (#408261) for 2 hours at 40°C, and amplified signals were detected using diaminobenzidine (#322310) or TSA Vivid™ Fluorophore 650 (#323273). Hematoxylin or DAPI (#ab104139, abcam) was used to stain nuclei. Images were captured using a Nikon Eclipse Ni microscope, the Nikon AXR inverted confocal microscope, or the Axio Scan Z7 (Zeiss).

### Ultrasonography

Descending aortas were imaged in vivo using a Vevo 3100 ultrasound system with a MicroScan MX550D transducer (25-55 MHz, FUJIFILM VisualSonics Inc) as described previously (27). Briefly, mice were anesthetized with isoflurane (1.0-2.5% vol/vol) adjusted to maintain a heart rate of 400-600 beats per minute during ultrasonography. Aortic luminal diameter was measured between the inner edge to inner edge of the vessel at both end systole and diastole. The β-index was determined according to the following formula (28, 29):

β-index = ln(Ps / Pd) / ((Ds − Dd) / Dd)

where Ps and Pd represent systolic and diastolic pressures, and Ds and Dd represent systolic and diastolic diameters, respectively. Since BAPN administration does not alter blood pressure (30), a blood pressure of 120/80 mmHg was assumed for calculation purposes.

### Biomechanical measurement

Descending aortic segments (distal to the left subclavian artery) were excised and cannulated at both ends with glass micropipettes, secured with 6-0 sutures, and mounted in a computer-controlled biomechanical testing system in Hanks’ buffered salt solution at room temperature to assess passive properties (31). Specimens were preconditioned by cyclic pressurization from 10 to 140 mmHg at the in vivo axial stretch. Mechanical testing included cyclic pressurization (10-140 mmHg) at three axial stretches (in vivo and ±5%) and cyclic axial loading at four fixed pressures (10, 60, 100, and 140 mmHg). Outer diameter, pressure, axial length, and axial force were continuously recorded, and biaxial mechanical properties were inferred using a validated four-fiber family constitutive model for the passive properties (32).

### Bulk RNA Sequencing

Descending thoracic aortas (Th2-Th6 region) were harvested from male control or BAPN-administered mice after 7, 28, and 84 days of administration (n = 4 per group). After removing periaortic tissues, aortic samples were incubated with RNAlater (#AM7020, Invitrogen) at 4°C overnight. Subsequently, RNA was extracted using RNeasy Fibrous Tissue Mini kits (#74704, Qiagen) and shipped to Novogene for bulk mRNA sequencing. Sequencing library was generated from total mRNA (1 µg) using NEBNext UltraTM RNA Library Prep Kits for Illumina (New England BioLabs). cDNA libraries were sequenced by a NovaSeq 6000 (Illumina) in a paired-end fashion. Adapter sequences were trimmed, and poly-N-containing reads and low-quality reads were removed using in-house Perl scripts by Novogene. Sequencing data in FASTQ format were aligned to the mouse reference genome assembly GRCm38 (mm10) using STAR (v2.5, mismatch=2) (33). Aligned reads were quantified using HTSeq (v0.6.1, -m union) (34). Raw read counts were normalized using the trimmed mean of M-values (TMM) method, and PCA was performed. Differentially expressed genes across groups were identified using edgeR. mRNA abundance of selected extracellular matrix (ECM)-related genes was visualized as a heatmap using pheatmap.

### Spatial Transcriptomic Analyses

Spatial transcriptomic analyses were performed as per Visium 2.0 kit guidelines (10x Genomics, Demonstrated Protocol CG000520, User Guide CG000495). Briefly, 5 µm sections of paraformaldehyde-fixed paraffin-embedded tissue blocks were mounted on microscope slides and dried for 3 hours at 42 °C. Dried sections were deparaffinized and stained with hematoxylin and eosin. Histological images were then captured on a motorized stage by a Nikon Ni-E microscope with a Nikon Digital Sight 10 CMOS camera. Image tiles were stitched with Nikon Elements software. Tissue on slides was immediately hybridized with the Visium Mouse Transcriptome Kit v2 (#PN-1000365, 10x Genomics). Ligated probes were captured on Visium slides using a CytAssist instrument (10x Genomics), and probe libraries were amplified. Paired-end sequencing was performed on an Illumina NovaX system in Advanced Genomics Core of the University of Michigan. Sequencing data were processed using Space Ranger HD software (10x Genomics) to generate gene expression matrices and spatial barcode information. Spots with fewer than 50 detected genes or fewer than 100 total counts were excluded. Downstream analyses were conducted using the Seurat R package (v4, Satija Lab) (35). Data were normalized using NormalizeData, and the top 3,000 variable features were identified using variance-stabilizing transformation. PCA was performed, followed by unsupervised clustering using a shared nearest-neighbor graph at a resolution of 2.0 and UMAP visualization based on the top 30 principal components. Cell clusters were annotated based on the expression of canonical marker genes for SMCs, endothelial cells, fibroblasts, and adipocytes. Differentially expressed genes (DEGs) between resident SMC and SMC-like cell clusters were identified using FindMarkers. Gene Ontology (GO) biological process analysis of genes enriched in SMC-like cells with Log_2_FC (fold change) > 0.25 was performed using clusterProfiler (36), with GO terms simplified using a similarity cutoff of 0.5. mRNA abundance of selected gene sets including SMC contractile, extracellular matrix, and TGF-β pathway genes was visualized as heatmaps using pheatmap, with expression values representing Log_2_FC relative to the mean across groups. Violin plots and spatial gene expression patterns were visualized using Seurat and ggplot2.

### In Vitro Experiments

*Descending aortas were* harvested and enzymatically digested using a cocktail containing collagenase type II (60 mg, #LS004176, Worthington), elastase (3.75 mg, #LS002292, Worthington), and DMEM (20 mL, #30-2002, ATCC) without fetal bovine serum but with penicillin-streptomycin (200 µL, #15070-063, Gibco). After an initial 10-minute incubation at 37°C, the adventitia was carefully peeled off and further digested for 60 minutes with gentle pipetting every 15 minutes. Sca-1 labeling was conducted directly in the cell suspension using Anti-Sca-1 (non-HSC) MicroBeads (10 µL, #130-106-641, Miltenyi Biotec), and MACS® buffer (1 mL, #130-091-376 and #130-091-221, Miltenyi Biotec) was then added to proceed with magnetic sorting. Labeled cells were first filtered through cell strainers (#10199-658, VWR), then separated using MS columns (#130-042-201, Miltenyi Biotec) mounted on a MiniMACS™ Separator (#130-042-102, Miltenyi Biotec) with a MACS® MultiStand (#130-042-303, Miltenyi Biotec). Sca-1 cells were collected by flushing the columns and subsequently centrifuged at 300 g for 10 minutes. The cells were then resuspended in culture medium and seeded onto 24-well plates (#10062-896, VWR) or chamber slides (#154534, Thermo Fisher Scientific). The culture medium (#30-2002, ATCC), including fetal bovine serum (10% vol/vol, #16140-071, Gibco) and penicillin-streptomycin (1% vol/vol, #30-2300, ATCC), was changed every two days. For qPCR, cells were dissolved in 300 μL of TRIzol™ Reagent (#15596018, Invitrogen), and RNA was extracted using Direct-zol RNA Microprep Kits (#R2060, Zymo Research).

### mRNA Quantification

RNA was extracted using the same protocol as that employed for bulk RNA sequencing. To quantify mRNA abundance, total RNA was reversely transcribed with an iScript™ cDNA Synthesis kit (#170-8891, Bio-Rad), and quantitative polymerase chain reaction (qPCR) was performed using Bio-Rad CFX384 real-time system with TaqMan Gene Expression Master Mix (#4369016, Thermo Fisher Scientific) and TaqMan Gene Expression Assays (Thermo Fisher Scientific) specific for each target gene (listed in the Major Resource Table). Data were analyzed using ΔΔCt method and normalized by the geometric mean of 3 reference genes (*Actb*, *Gapdh*, and *Ppia*). For pre-mRNA quantification, one-step RT-qPCR was performed directly from total RNA using iTaq™ Universal SYBR® Green One-Step Kit (#172-5150, Bio-Rad) and intron-targeting primers.

### Transmission Electron Microscopy

Descending aortas were fixed in glutaraldehyde (2.5% wt/vol) in sodium cacodylate buffer (0.1 M, pH 7.4, #15960-01, Electron Microscopy Sciences) and stored at 4°C. Samples were post-fixed in osmium tetroxide (1% wt/vol, #19150, Electron Microscopy Sciences) and potassium ferrocyanide (1.5% wt/vol, #152560, MP Biomedicals) for 1 hour at room temperature, followed by staining with uranyl acetate (1% wt/vol, #22400, Electron Microscopy Sciences) at 4°C for 1 hour. After dehydration by propylene oxide (#20401, Electron Microscopy Sciences), samples were infiltrated overnight with a 1:1 mixture of propylene oxide and EMbed 812 resin (#14900, Electron Microscopy Sciences), mixed with Dodecenyl Succinic Anhydride (DDSA), Methyl-5-Norbornene-2,3-Dicarboxylic Anhydride (NMA), and 2,4,6-Tris(dimethylaminomethyl)phenol (DMP-30) (#14900, #13710, #19000, #13600, Electron Microscopy Sciences). Ultrathin sections (60 nm) were cut using a diamond knife and mounted on cupper grids (#01802-F, TED PELLA, INC.). Images were captured using Talos F200X (Thermo Fisher Scientific).

### Statistics

SigmaPlot 15 (SYSTAT Software Inc.) was used for statistical analyses. Data are presented as box-and-whisker plots showing the median and 25^th^-75th percentiles. Normality and homogeneity of variance assumptions were assessed by Shapiro-Wilk and Brown-Forsythe tests, respectively. For comparisons between two groups, Student’s t-test (for normally distributed data with equal variance), Welch’s t-test (for normally distributed data with unequal variance), or Mann-Whitney U test (for non-normally distributed data) was used. Because both normality and homogeneity of variance assumptions were satisfied for comparisons among three or more groups, one-way ANOVA followed by the Holm-Sidak post hoc test was performed. For repeated measures from the same animals, repeated-measures ANOVA followed by Bonferroni’s post hoc test was used. Survival curves were generated using the Kaplan-Meier method and compared using the log-rank test. P or False discovery rate (FDR)-adjusted P values < 0.05 was considered statistically significant.

### Study Approval

For human samples, all patients provided written informed consent upon enrollment. This study was conducted in accordance with the guidelines of the Declaration of Helsinki and was approved by the Ethics Committee of Kobe University (#B210014). All mouse experiments were conducted in accordance with the ARRIVE guidelines (Animal Research: Reporting of In Vivo Experiments) and were performed with the approval of the University of Kentucky Institutional Animal Care and Use Committee (University of Kentucky IACUC protocol number: 2018-2967).

### Data Availability

All numerical data are available in Supplemental Excel File. All raw data, including bulk RNA sequencing and spatial transcriptomic data, are available upon reasonable request to the corresponding author.

## RESULTS

### Elastic fibers were synthesized in the vascular wall of the false lumen in patients with Stanford type B AD

We first investigated whether the synthesis of elastic fibers occurs in patients with AD. Aortic samples were obtained at 3 weeks, 6 months, and 15 years after diagnosis of the onset of Stanford type B AD (**Figure 1A**). Patients’ characteristics are summarized in **Table 1**. AD was diagnosed based on contrast-enhanced computed tomography demonstrating false lumen formation. The onset of AD was defined as the time of presentation with acute back or chest pain. Verhoeff’s iron hematoxylin staining demonstrated that, in the samples obtained 3 weeks after onset, pale purple layers were distributed on the surface of the false lumen wall. Higher magnification revealed that these layers were composed of thin and short elastic fibers. In the sample collected 6 months after onset, distinct elastic fibers were evident on the surface of the false lumen wall. These fibers were also observed in the wall of the false lumen at 15 years after onset (**Figure 1B**). In situ hybridization revealed that elastin mRNA was abundant in regions showing de novo elastic fiber formation in the false lumen wall at 3 weeks after the onset, although it was reduced in ADs obtained at 6 months and 15 years after onset (**Figure 1C**). Immunostaining demonstrated co-localization of elastin mRNA with myosin heavy chain 11 (MYH11), a SMC marker (**Figure 1D**). Taken together, these data suggest that, even during adulthood, the aortic wall retains the capacity to resynthesize elastic fibers, a process that is reactivated in cells with an SMC phenotype following AD.

**Figure 1.**
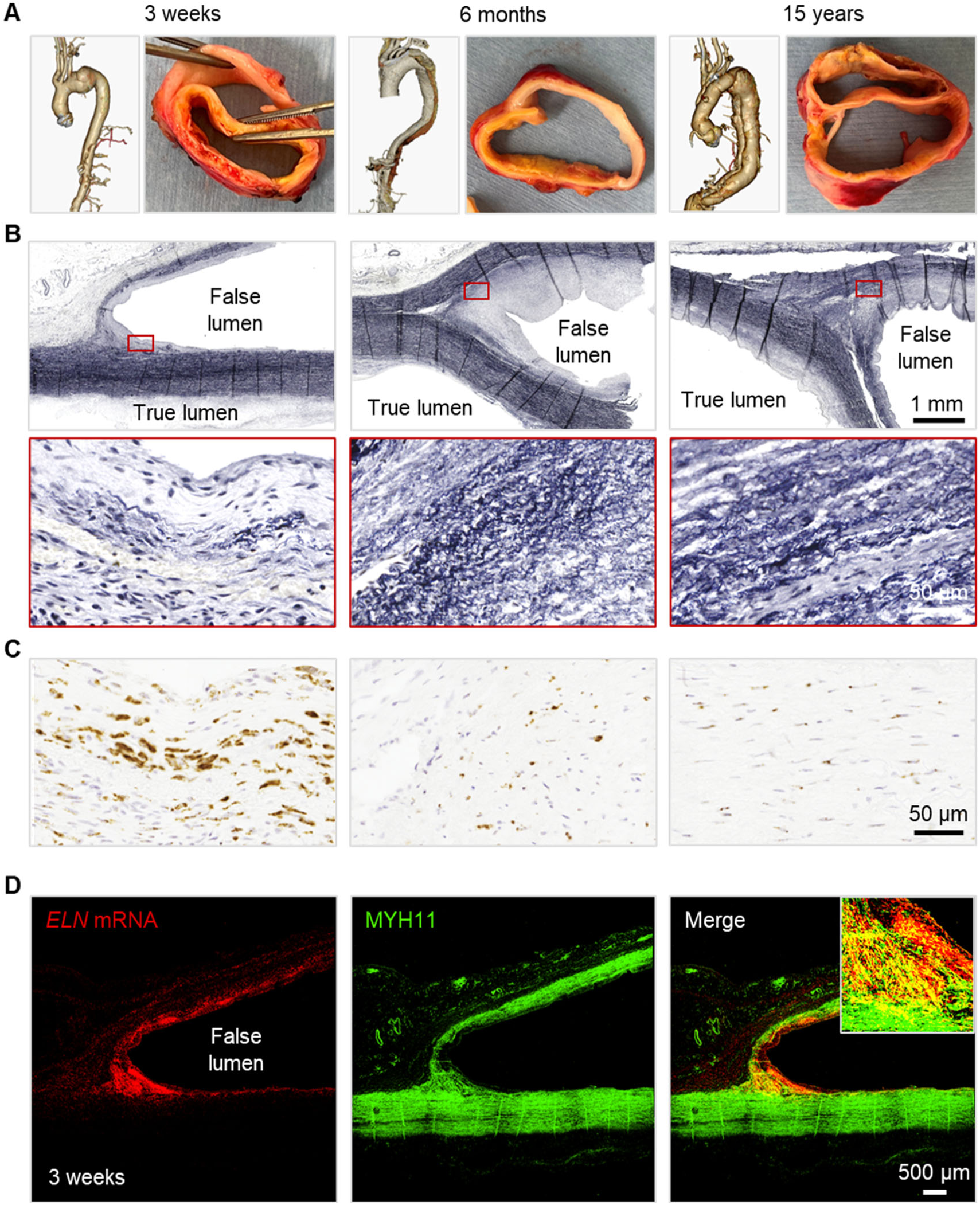
De novo elastic fiber formation following Stanford type B AD in humans. Representative **(A)** 3D CT and gross images, **(B)** Verhoeff’s iron hematoxylin, and **(C)** in situ hybridization for *ELN* mRNA at 3 weeks, 6 months, and 15 years after the onset of Stanford type B AD. High-magnification images were captured at red boxes. **(D)** Representative images of immunostaining for MYH11 combined with *ELN* mRNA in situ hybridization at 3 weeks after AD onset.

**Table 1.**
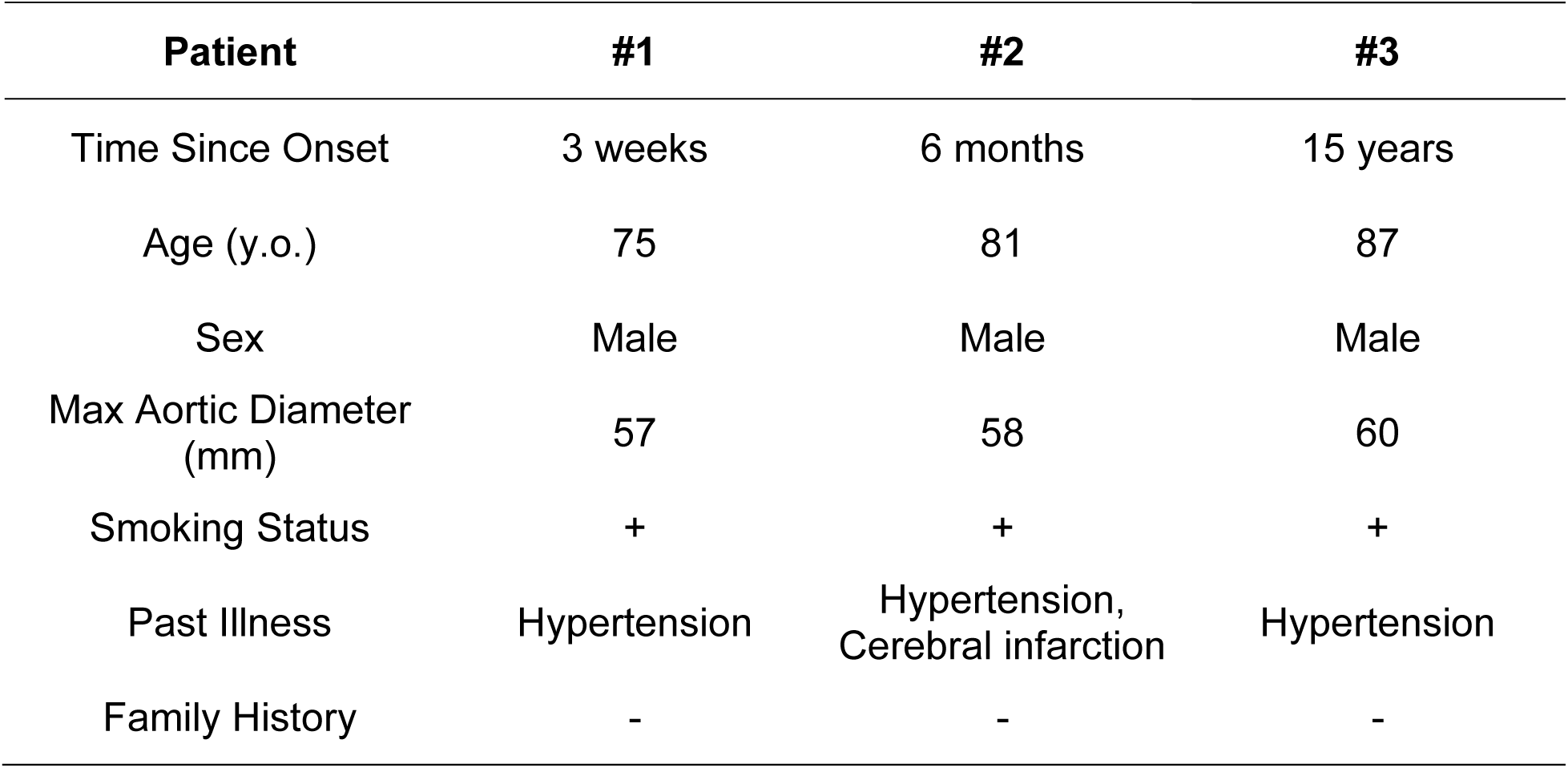
Patients’ Characteristics of Human AD Samples.

### De novo elastic fibers were synthesized in the false lumen wall in chronic AD of BAPN-administered mice

We investigated whether de novo elastic fiber synthesis occurs in AD in mice following BAPN administration. Our previous study demonstrated that oral administration of BAPN recapitulates multiple facets of human Stanford type B AD, including medial disruption, ECM degradation, and AD lesion localized in the descending aorta. Most mice administered BAPN for 1 week showed no overt aortic pathology, representing a pre-AD stage. During the first 4 weeks of BAPN administration, approximately 25% of mice died from aortic rupture, increasing to nearly 50% by 12 weeks (**Figure 2A**). By 4 weeks, mice developed acute AD characterized by fresh thrombus in the media of the descending aorta, and by 12 weeks there was false lumen enlargement with vascular wall thickening, which is considered to represent chronic AD (**Figure 2B, C**). Bulk RNA sequencing was performed using descending aortic tissues collected from vehicle- and BAPN-administered mice at the pre AD (1 week), acute AD (4 weeks), and chronic AD (12 weeks) stages. Principal component analysis (PCA) revealed distinct transcriptomic profiles in acute and chronic phases of AD (**Figure 2D**). Consistent with these alterations, aortic elastin (*Eln*) mRNA abundance was not increased during either pre or acute phases but was elevated significantly during the chronic phase. In addition to *Eln*, elastin-related genes, such as *Fbn1*, *Fbln5*, and *Lox*, were also upregulated in chronic AD (**Figure 2E**). qPCR validated the increase of *Eln* and elastin-related gene mRNA abundance in chronic, but not pre or acute, AD of BAPN-administered mice compared to vehicle controls (**Figure 2F**, **Supplemental Figure 1**). Since *Eln* mRNA expression is also regulated by post-transcriptional modification of pre-mRNA (37–40), *Eln* pre-mRNA was quantified by a one-step RT-qPCR method using primers targeting intron 21 of *Eln* gene. Consistent with *Eln* mRNA, *Eln* pre-mRNA abundance was increased significantly in chronic AD (**Figure 2G**). These results provide compelling evidence that *Eln* transcription is upregulated at a protracted interval after AD.

**Figure 2.**
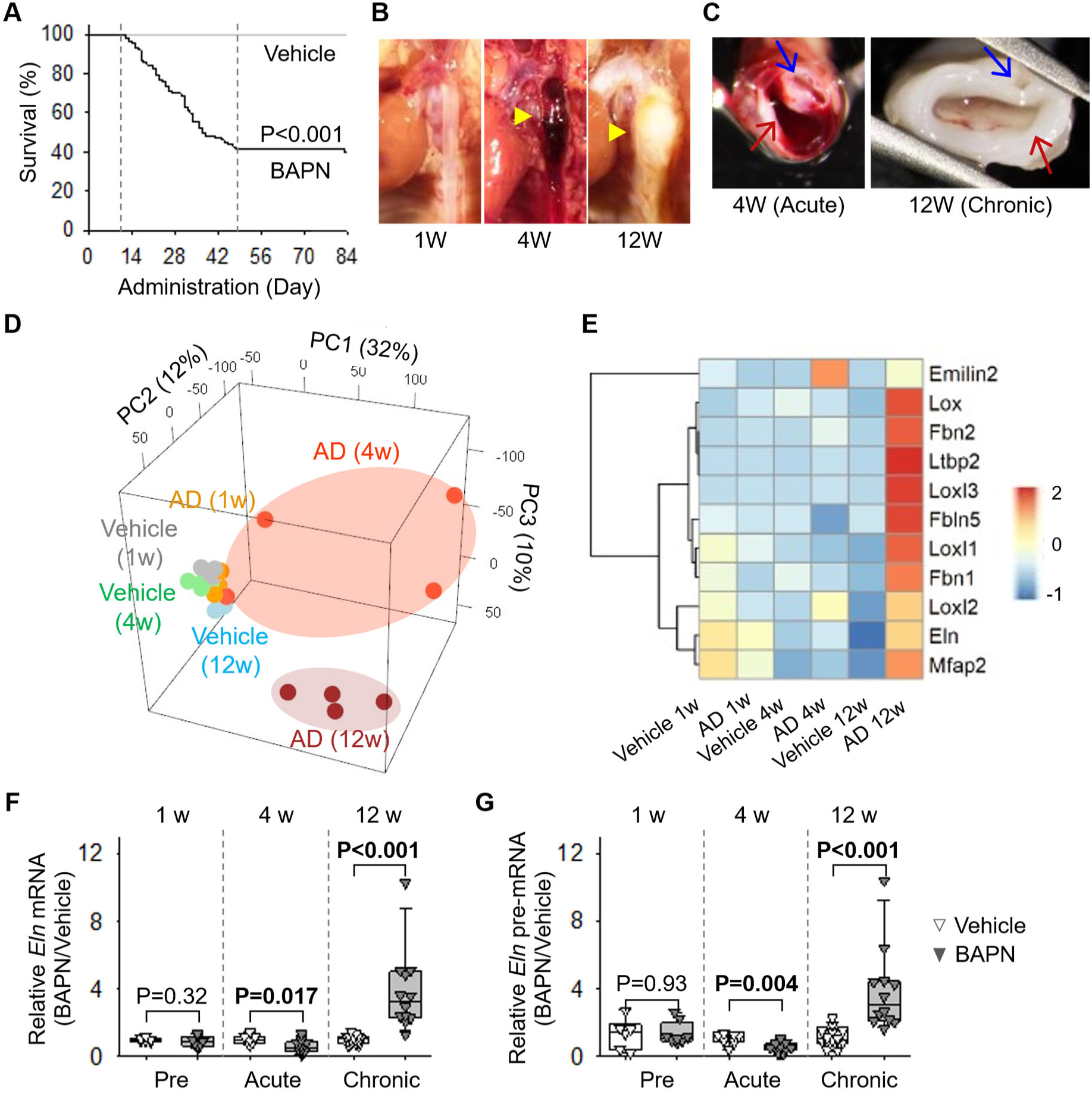
Reactivation of elastin and elastic fiber-related mRNA transcription following AD in mice. **(A)** Survival curve of BAPN-administered mice. N=46 and 168 for vehicle and BAPN, respectively. P value was determined by Log-Rank test. **(B)** Representative in situ images of AD at pre, acute, and chronic phases. **(C)** Cross-sectional ex vivo images of AD at acute and chronic phases. **(D)** Principal component analysis (PCA) plot and **(E)** heatmap of elastic fiber-related genes in bulk RNA sequencing using aortic tissues at the pre, acute, and chronic phases of BAPN-induced AD and vehicle controls at each interval. qPCR for **(F)** *Eln* mRNA and **(G)** *Eln* pre-mRNA abundance at pre, acute, and chronic phases following BAPN-induced AD and vehicle controls. N=6-15/group. P values were determined by Welch’s t-test or Mann-Whitney test between administrations at each time point.

To determine the spatial distribution of *Eln* mRNA, *in situ* hybridization was performed. *Eln* mRNA was distributed mainly in the false lumen wall (**Figure 3A**). Immunostaining demonstrated that elastin protein was also present in the false lumen wall, largely overlapping with *Eln* mRNA (**Figure 3B**). As observed in human ADs, numerous elastic fibers were observed in the vascular wall of the false lumen in Verhoeff’s iron hematoxylin staining (**Figure 3C**). It is worth noting that de novo elastic fiber formation was not observed in mice that died of false lumen rupture in the chronic phase (**Supplemental Figure 2**). These findings suggest that newly formed elastic fibers may exert a protective effect. However, compared to native elastic fibers around the true lumen, de novo elastic fibers in the false lumen wall showed a distinct structure. New fibers had more layers, but were more densely packed, thinner, and less organized (**Figure 3D**). The structure of de novo fibers was further evaluated by 3-dimensional confocal microscopy. Z-stack confocal images with staining for Alexa Fluor 633 hydrazide, a specific dye for elastic fibers, revealed that native fibers circumscribing the true lumen were thick and displayed a waved structure, whereas de novo fibers showed a spider web-like structure (**Figure 3E**). Transmission electron microscopy demonstrated that native elastic fibers had smooth surfaces, whereas de novo elastic fibers were composed of small granules (**Figure 3F**).

**Figure 3.**
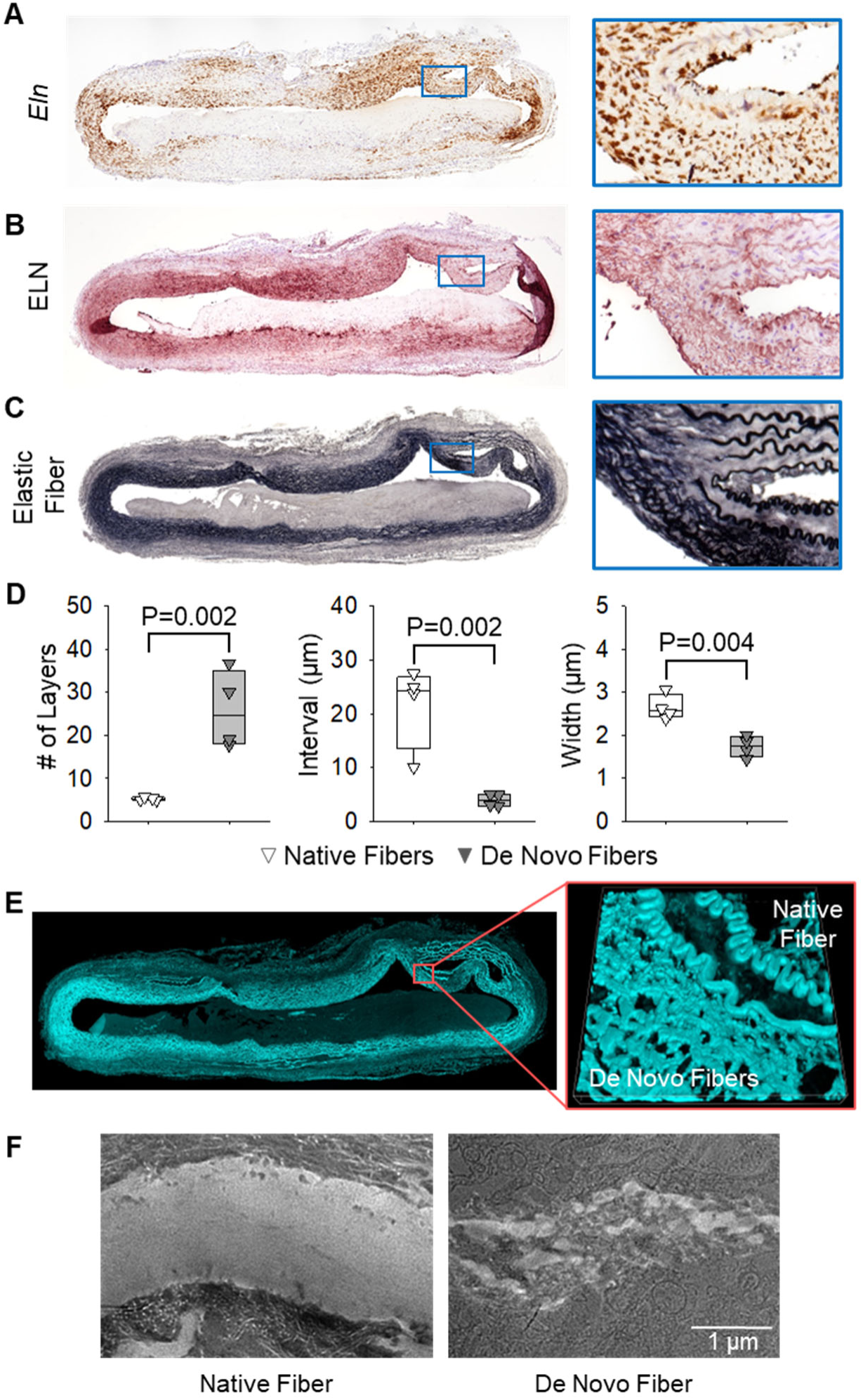
De novo elastic fiber formation in the vascular wall of the false lumen in AD at the chronic phase. Representative images of **(A)** in situ hybridization for *Eln* mRNA, **(B)** immunostaining for elastin, and **(C)** Verhoeff’s iron hematoxylin staining in BAPN-induced AD at the chronic phase. **(D)** Quantification of the number of elastic layers, layer intervals, and layer width. N=4/group. P values were determined by Welch’s t-test or Mann-Whitney U-test. **(E)** Representative images of Alexa 633 elastic fiber staining of the chronic AD sample from BAPN-administered mice. **(F)** Representative images of transmission electron microscopy of native and de novo elastic fibers in chronic AD of BAPN-administered mice.

### Circumferential stiffness was increased in chronic AD of BAPN-administered mice

Since de novo elastic fibers exhibited structural differences from native fibers, we next examined biomechanical properties of AD samples. Ultrasonography was performed in control and AD mice at pre, acute, and chronic phases, and vascular wall elasticity and stiffness were calculated. Although we used a high-frequency ultrasound system using a 55 MHz transducer, the spatial resolution was limited to measure accurate wall thickness in mice, which is needed to calculate vascular elasticity. Therefore, reliable elasticity data could not be obtained. In contrast, structural stiffness, which is primarily based on diameter measurements, was able to be assessed. Compared to control aortas, β-index, an index of structural stiffness (28, 29), was elevated significantly in dissected aortas over time (**Figure 4A**). To validate these findings and measure vascular elasticity, we also evaluated biomechanical properties using an ex vivo system that enabled precise measurements of pressure-diameter relationships and calculation of mechanical parameters under physiological loading conditions. The descending thoracic aorta was cannulated and subjected to controlled intraluminal pressure (**Figure 4B**). Stored elastic energy was reduced in chronic AD, indicating a loss of elastic fiber functionality. Axial stiffness was decreased, whereas circumferential stiffness was increased, reflecting anisotropic mechanical remodeling. Distensibility was also reduced, consistent with the in vivo measurements of the β-index (**Figure 4C**). Pressure-diameter curves demonstrated reduced expansion of the aortic wall in the chronic phase of AD in response to pressure (**Figure 4D**). Collectively, these findings indicate that the vascular wall of chronic AD is stiffer than the normal aortic wall.

**Figure 4.**
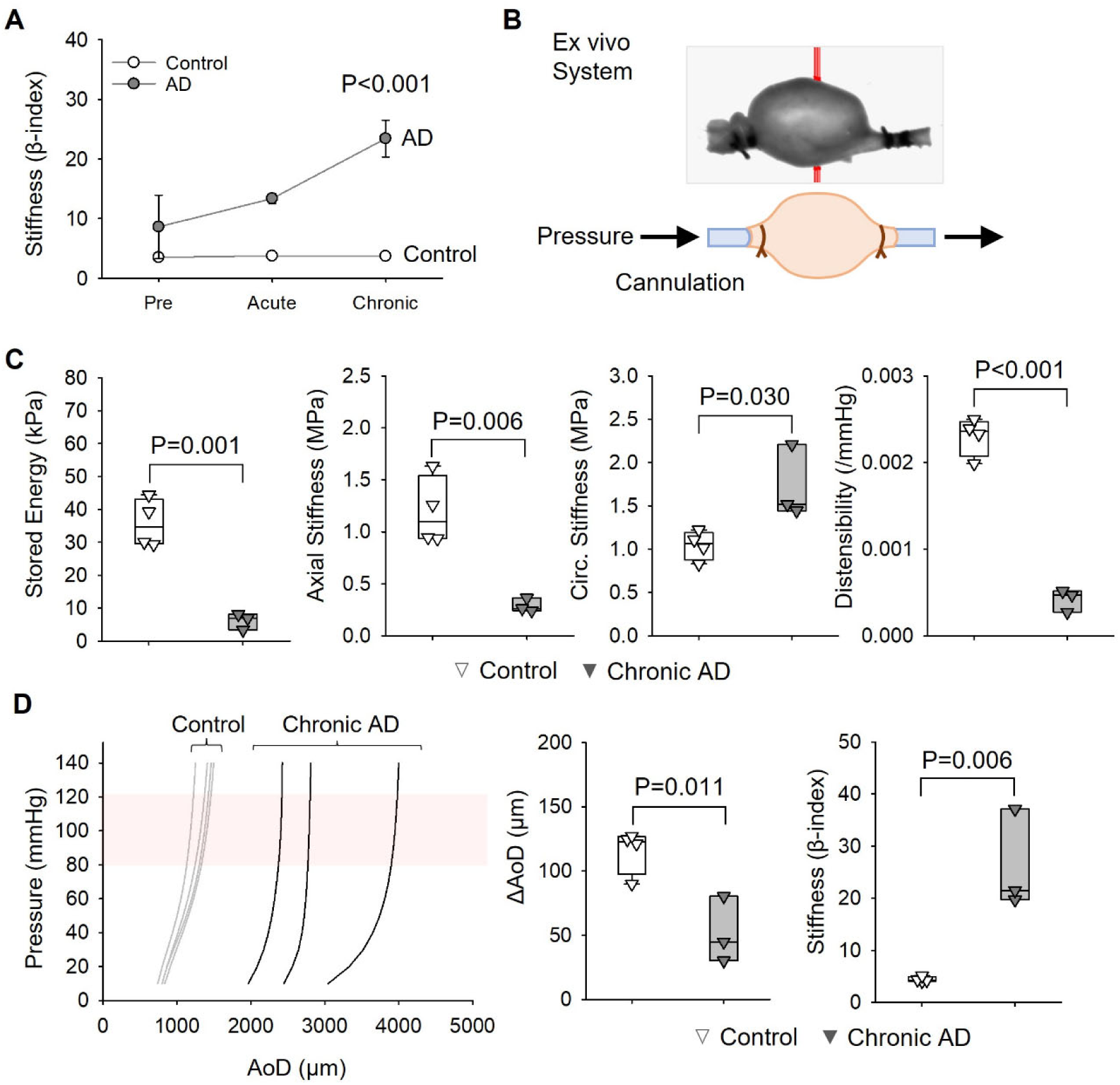
Increase in vascular stiffness in chronic AD of BAPN-administered mice. **(A)** Vascular wall stiffness measured by ultrasonography at the pre-, acute-, and chronic phases following BAPN-induced AD. N=3-4/group. P values were determined by repeated measures ANOVA followed by Bonferroni t-test. **(B)** Schematic image of the ex vivo biaxial mechanical testing apparatus for aortic biomechanical measurements. **(C)** Quantification of aortic mechanical properties in vehicle control and chronic BAPN-induced AD samples. N=3-4/group. Stored energy and axial and circumferential (Circ.) stiffness were measured at 100 mmHg. Distensibility is calculated over the physiological pressure range of 80-120 mmHg. **(D)** Pressure-diameter curves, the change of aortic diameters (AoD) between 80 and 120 mmHg, and β-index wall stiffness. P values were determined by Student’s t-test.

### SMC-like cells were the source of de novo elastic fibers in chronic AD of BAPN-administered mice

To identify the cell types responsible for de novo elastic fiber formation, immunostaining for aortic cell markers was performed in chronic AD of BAPN-administered mice. The lumen of the vascular wall of the false lumen was partially covered by CD31-positive endothelial cells, whereas the outer layer was covered by ER-TR7-positive cells as a marker of fibroblasts (**Figure 5A, B**). Both MYH11 and α-smooth muscle actin (αSMA) were extensively distributed in the false lumen wall (**Figure 5C, D**). Co-staining of αSMA and MYH11 with in situ hybridization of *Eln* mRNA demonstrated that more than 60% of *Eln* mRNA-positive cells were colocalized with αSMA and MYH11-positive cells in the false lumen wall (**Figure 5E, F**). These data indicate that the false lumen wall was primarily composed of cells expressing SMC markers and that these cells contribute to de novo elastic fiber formation in chronic AD.

**Figure 5.**
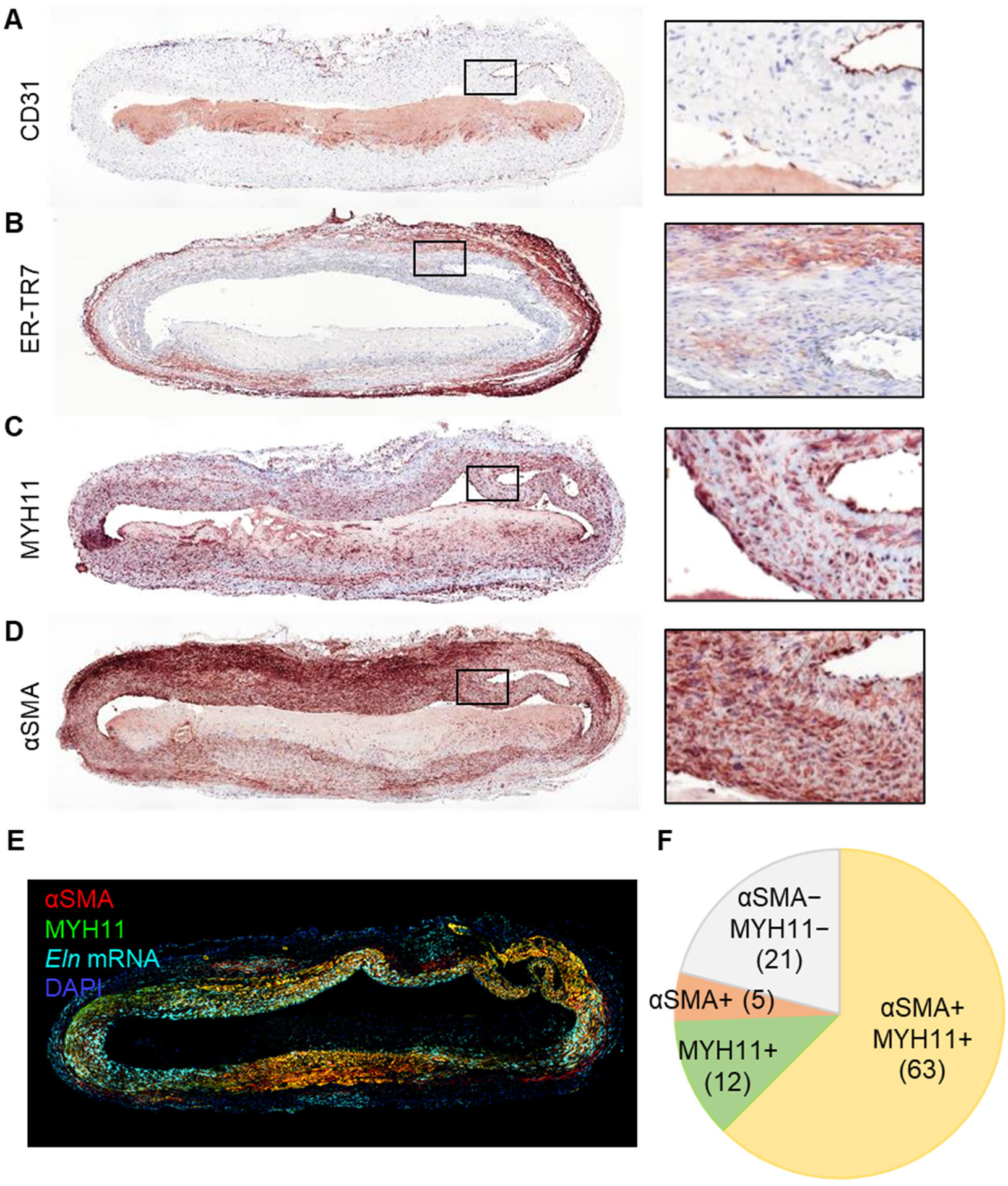
Expansion of cells expressing SMC markers in the vascular wall of the false lumen in chronic AD. Representative images of immunostaining for **(A)** CD31, **(B)** ER-TR7, **(C)** MYH11, **(D)** αSMA, and **(E)** confocal image of double-immunostaining for αSMA and MYH11 combined with in situ hybridization for *Eln* mRNA in chronic AD of BAPN-administered mouse. **(F)** Proportion of SMC-marker-positivity in cells expressing *Eln* mRNA.

Expansion of cells expressing SMC markers in the false lumen wall prompted us to investigate their clonality as a potential mechanism underlying de novo elastic fiber synthesis following AD. Lineage tracing that targeted resident SMCs was performed using *Acta2-*CreER^T2^ mice with the “Rainbow” reporter. Rainbow mice have four fluorescent proteins, eGFP, mOrange, mCerulean, and mCherry, in the *Rosa26* locus (**Figure 6A**) (41–43). eGFP is expressed in all cells at baseline. Once *Acta2*-CreER^T2^ is translocated to the nuclei, cells in which the *Acta2* promoter is active (encoding αSMA), the cell color change from eGFP to mOrange, mCerulean, or mCherry, with random and equal distribution. Thus, polyclonal proliferation of SMCs in the false lumen wall would be indicated by the presence of multiple colors, with red, orange, and light blue. In contrast, monoclonal proliferation would be detected by the presence of one single recombinant color, except for green. To track SMC-lineage cells, tamoxifen was injected before AD induction by BAPN administration. Descending AD was harvested after 12 weeks of BAPN administration (**Figure 6A**). Confocal imaging revealed that medial SMCs surrounding the true lumen of the descending aorta exhibited rainbow colors, indicating the system was functional (**Figure 6B**). Surprisingly, while partially composed of a single recombinant color, the false lumen wall was primarily composed of cells expressing eGFP (**Figure 6B**). Meanwhile, these cells expressed both MYH11 and αSMA in immunostaining (**Figure 6C**), indicating their SMC phenotype. Since DNA recombination by tamoxifen injection was induced before AD formation, resident SMCs and their progeny expressed mOrange, mCerulean, or mCherry; whereas other cell types expressed eGFP. Therefore, these results suggest that cells expressing SMC markers in the false lumen wall were not derived from resident SMCs in the aortic media. Instead, non-SMC lineage cells differentiated into SMC-like cells after AD induction.

**Figure 6.**
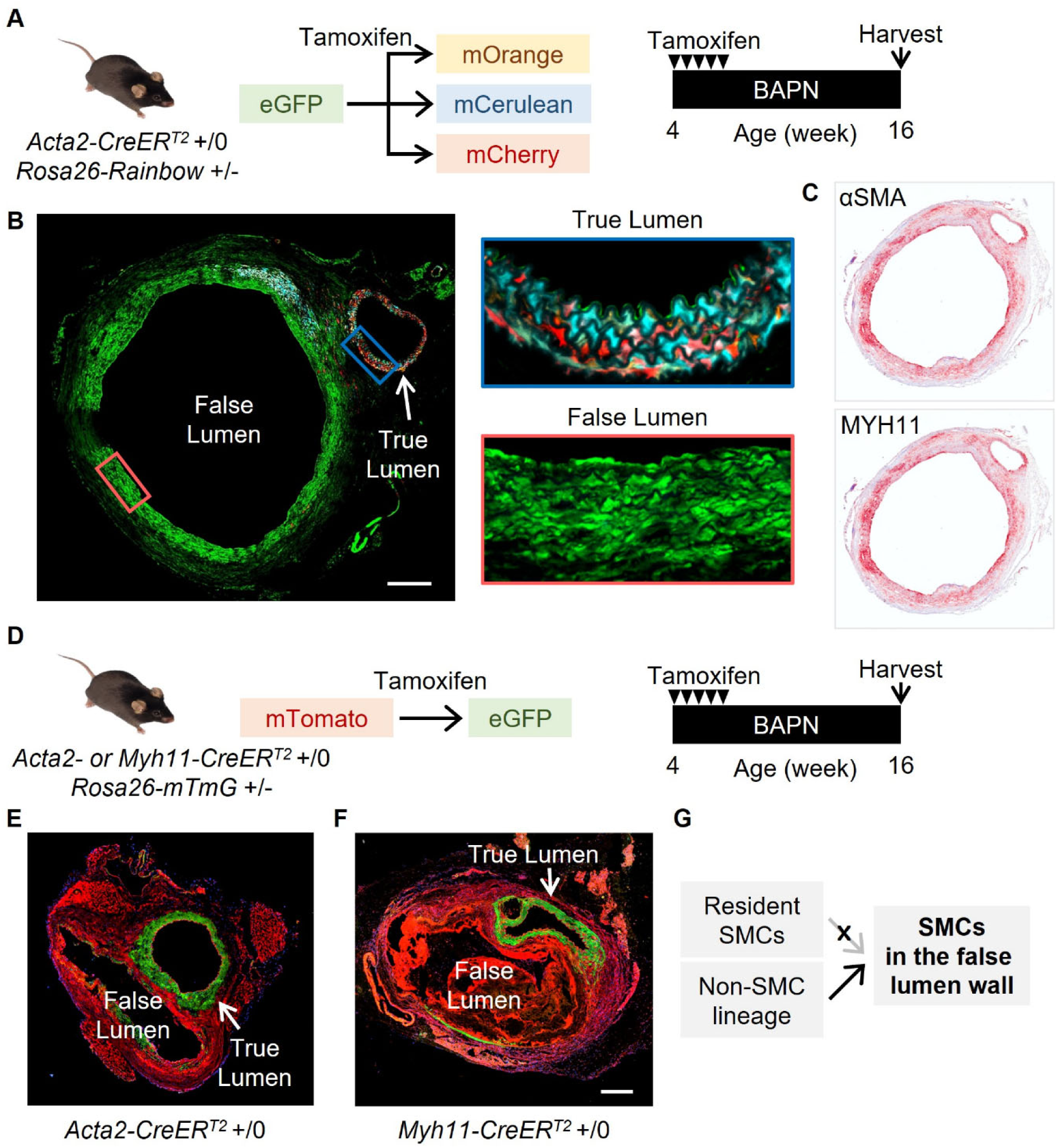
Lineage tracings for SMCs in the false lumen wall of BAPN-induced chronic AD. **(A)** Experimental design of Cre-mediated multicolor lineage tracing using “Rainbow” mice with tamoxifen injection and BAPN administration. Representative images of **(B)** lineage tracing and **(C)** immunostaining for MYH11 and αSMA in BAPN-induced chronic AD from *Acta2*-CreER^T2^; *Rosa26*-Rainbow mice. **(D)** Schematic of binary Cre-mediated lineage labeling using “mTmG” reporter mice and the experimental strategy for tamoxifen injection and BAPN administration. Representative confocal images of chronic AD in **(E)** *Acta2*- and **(F)** *Myh11-*CreER^T2^*; Rosa26-*mTmG mice. **(G)** Diagram illustrating the potential cellular sources of cells expressing SMC markers in the false lumen of chronic AD. Scale bar = 250 µm.

To further validate and strengthen these findings, additional lineage tracing was performed using another reporter system. *Acta2*-CreER^T2^ and *Myh11*-CreER^T2^ mice were bred with a *Rosa26-mTmG* reporter line to track SMC lineage cells (**Figure 6D**). In this system, cells constitutively express membrane-targeted tdTomato (mTomato, red) at baseline, whereas Cre-mediated recombination switches expression to membrane-targeted GFP (mGFP, green), resulting in a red-to-green color conversion (44). Tamoxifen was injected to label resident SMCs at the same time as BAPN initiation, thereby inducing recombination in resident SMCs. Aortic tissues were harvested 12 weeks after BAPN administration (**Figure 6D**). As expected, cells in the true lumen wall were predominantly GFP-positive, confirming efficient labeling of resident medial SMCs. In contrast, the false lumen wall remained largely tdTomato-positive (red), indicating the absence of Cre recombination in these cells (**Figure 6E, F**). Thus, these results further validate the differentiation of non-SMC lineage cells into SMC-like cells following AD (**Figure 6G**).

### Spatial transcriptomic analyses revealed distinct transcriptomic profiles between resident SMCs and SMC-like cells that drive de novo elastic fiber synthesis in the false lumen wall

Spatial transcriptomic analysis enables direct comparison of cellular transcriptomes located across spatially distinct regions. Therefore, to understand the difference in molecular signatures between resident SMCs (rSMCs) and elastogenic SMC-like cells, we performed spatial transcriptomic analysis using a chronic-AD sample of BAPN-administered mice (**Figure 7A**). In an unsupervised clustering analysis, SMC-like cells from the false lumen formed a distinct cluster, separate from rSMCs around the true lumen (**Figure 7B, C**). Consistent with the histological findings, spatial feature plots showed the colocalization of *Eln* mRNA with *Myh11* and *Acta2* mRNA in the false lumen wall (**Figure 7D**). To further characterize elastogenic SMC-like cells, SMC-like cells and rSMCs were extracted selectively and performed comparative analyses (**Figure 7E**). Consistent with the global UMAP, SMC-like cells and rSMCs formed distinct clusters (**Figure 7F**). Gene ontology analysis of genes significantly upregulated in SMC-like cells relative to rSMCs revealed enrichment of pathways associated with ECM organization, cell-substrate adhesion, and TGF-β signaling (**Figure 7G**). Heatmap analysis demonstrated higher abundance of ECM-related genes, including elastin and multiple collagen isoforms, in SMC-like cells. Genes associated with TGF-β signaling showed distinct transcriptional profiles in SMC-like cells. In addition, SMC-like cells exhibited relatively reduced abundance of canonical SMC contractile markers (**Figure 7H**). Violin plots also demonstrated significantly higher *Eln* mRNA abundance in SMC-like cells compared to rSMCs (**Figure 7I**). In contrast, contractile SMC markers exhibited lower abundance in SMC-like cells (**Figure 7J**), indicating that SMC-like cells represent an immature stage compared to rSMCs. Since immature SMCs may reflect recent differentiation, and prior studies have shown that adventitial progenitor cells can give rise to SMC-like cells (45–47), we focused on adventitial progenitor features. Consistent with this concept, several progenitor marker genes were enriched in SMC-like cells (**Figure 7H**). Of note, feature plot analysis revealed enrichment of *CD34*, a transmembrane glycoprotein reported previously as a marker for adventitial progenitor cells in the aorta (47–49), mRNA in the false lumen wall. In addition to *CD34*, other markers associated with adventitial progenitor cells, such as *Thy1* and *Aldh1a1* (50, 51), were also detected in SMC-like cells of the false lumen wall (**Figure 7K**). These findings suggest that, following AD, adventitial progenitor cells differentiate into SMC-like cells and acquire the capacity for de novo elastic fiber synthesis.

**Figure 7.**
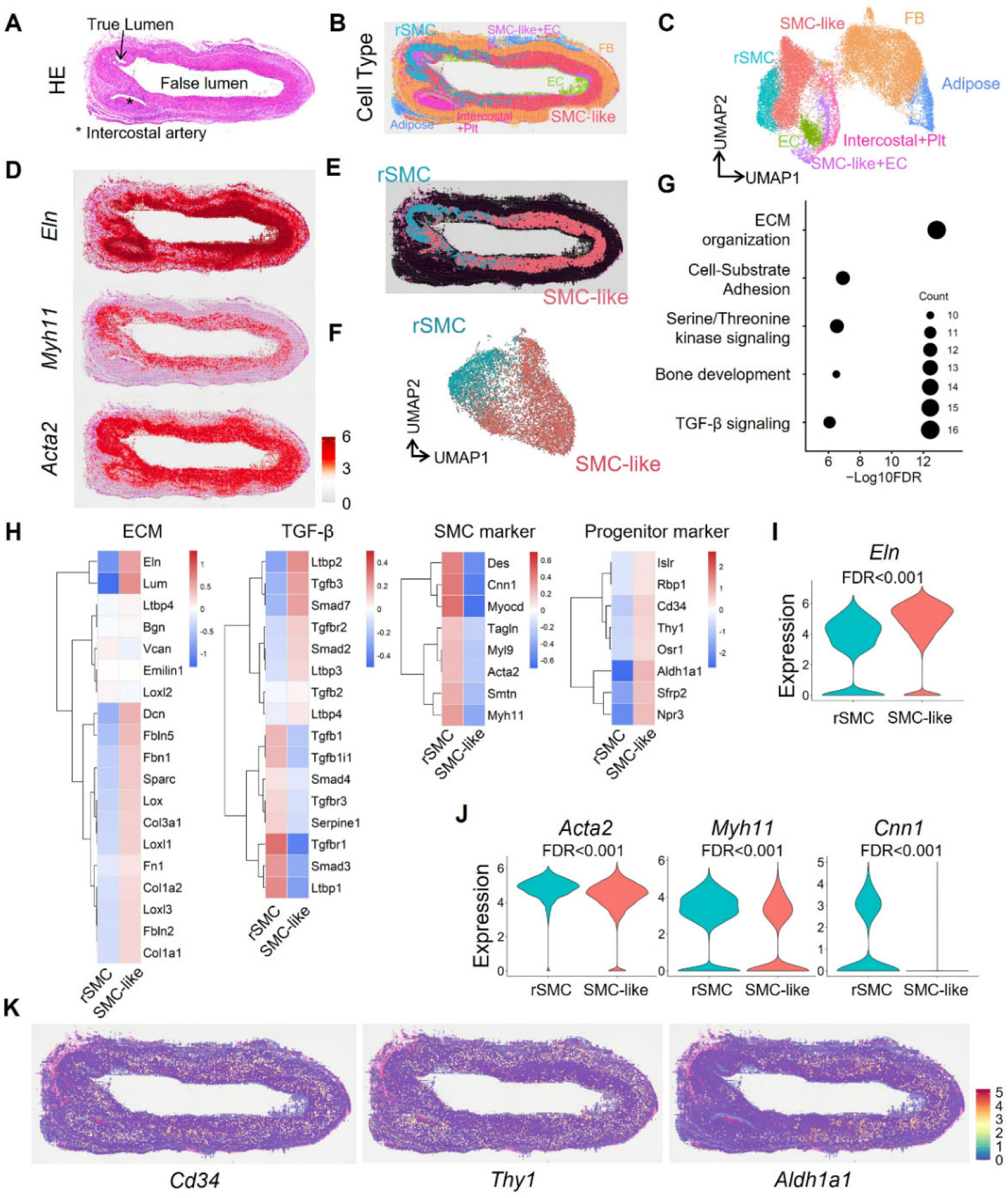
Cellular heterogeneity and the characteristics of SMC-like cells in the false lumen wall of chronic AD. **(A)** Representative hematoxylin and eosin (HE) staining of chronic AD in BAPN-administered mice. **(B)** Spatial annotation of major vascular cell types. **(C)** UMAP for aortic cells. **(D)** Spatial feature plots of *Eln*, *Myh11*, and *Acta2* mRNA in chronic AD. **(E)** Spatial projection of resident SMC (rSMC) and SMC-like cells within chronic AD. **(F)** UMAP for rSMC and SMC-like cells. **(G)** Gene ontology enrichment analysis of differentially expressed genes (DEGs) between rSMCs and SMC-like cells. **(H)** Heatmaps for major ECM, TGF-β, SMC marker, and progenitor marker genes. Violin plots for **(I)** *Eln* and **(J)** *Acta2*, *Myh11*, and *Cnn1* mRNA. Statistical significance was determined by Wilcoxon rank-sum test. FDR was calculated using the Benjamini-Hochberg method. **(K)** Spatial distribution of *Cd34*, *Thy1*, and *Aldh1a1* in BAPN-induced chronic AD.

### Adventitial progenitor cells were expanded in the false lumen wall following AD

To determine whether adventitial progenitor cells are involved in the expansion of SMC-like cells following AD, immunostaining for progenitor cell markers was performed in chronic AD of BAPN-administered mice. CD34 was detected widely within the false lumen wall (**Figure 8A**). Stem cell antigen 1 (Sca1), another marker for progenitor cells (43, 45–48, 52–57), was also broadly distributed throughout the false lumen wall of chronic AD (**Figure 8B**), in contrast to its predominantly adventitial localization at baseline and the acute phase (**Supplemental Figure 3**). Triple immunostaining for Sca1, αSMA, and MYH11 identified selected SMC-like cells expressing Sca1 in the false lumen wall (**Figure 8C**). Since Sca1 is also expressed in hematopoietic and endothelial cells, the presence of Sca1-positive cells does not necessarily indicate adventitial progenitor cells. Therefore, we then performed lineage tracing using *Gli1*-CreER^T2^ mice (**Figure 8D**). Gli1 is a Hedgehog-responsive transcription factor that specifically labels adventitial progenitor cells (56). At baseline, Gli1-lineage cells resided only around the aortic adventitia (**Supplemental Figure 4**), whereas those cells expanded within the false lumen wall of chronic AD (**Figure 8E**). These data are compelling evidence that adventitial progenitor cells differentiate into SMC-like cells in the false lumen wall during the chronic phase of AD.

**Figure 8.**
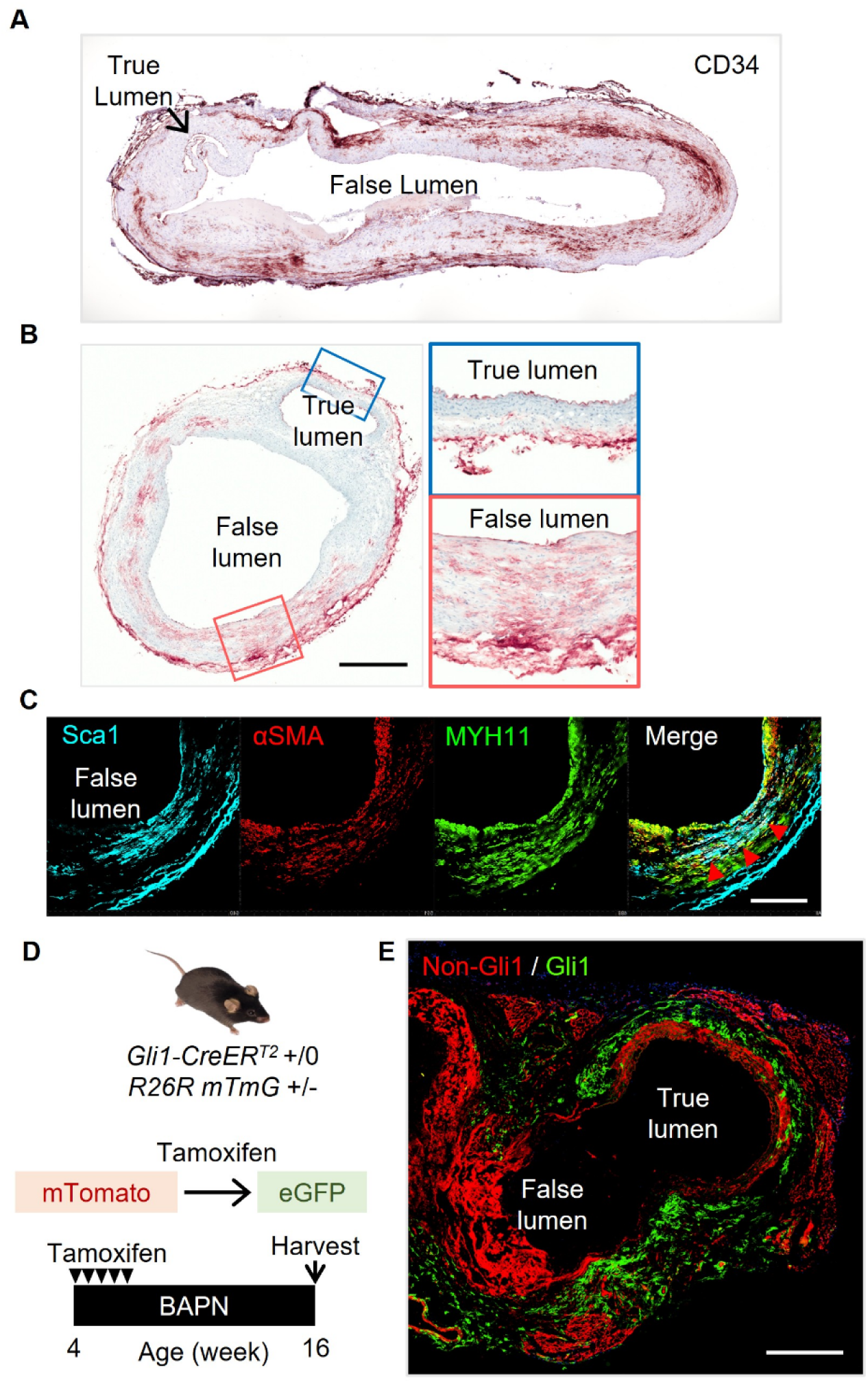
Distribution and expansion of vascular progenitor cells in the false lumen wall in chronic AD. **(A)** Representative image of immunostaining for CD34 in BAPN-induced chronic AD. Representative images of **(B)** immunostaining for Sca1 (*Ly6a*) in control, acute AD, and chronic AD, and **(C)** triple immunofluorescent staining for Sca1, αSMA, and MYH11 in chronic AD. **(D)** Schematic of the experimental strategy for lineage tracing using *Gli1*-CreER^T2^; *Rosa26*-mTmG mice. **(E)** Representative image of lineage tracing of Gli1-lineage cells in BAPN-induced chronic AD. Scale bar = 250 (A-C) or 500 (E) µm.

### Adventitial progenitor cells differentiated into elastogenic SMC-like cells

Finally, we performed in vitro experiments to investigate whether SMC-like cells derived from adventitial progenitor cells contribute to de novo elastic fiber synthesis. Descending aortas were harvested from C57BL/6J mice, and the adventitial layer was isolated by enzymatic digestion. Subsequently, adventitial cells were dissociated using multiple enzymes, and adventitial progenitor cells were selectively isolated using antibody-conjugated magnetic beads, followed by in vitro culture (**Figure 9A**). After 14 days of culture, mRNA abundance of SMC markers, *Acta2* and *Myh11,* was significantly increased (**Figure 9B**). Importantly, *Eln* mRNA was also increased in these SMC-like cells (**Figure 9B**). Immunofluorescent staining demonstrated that these cells expressed MYH11 after 14-day culture (**Figure 9C**). Of note, multiple elastic fibers were observed in immunofluorescent staining for elastin (**Figure 9D**). These data indicate that adventitial progenitor cells differentiate into SMC-like cells and form de novo elastic fibers.

**Figure 9.**
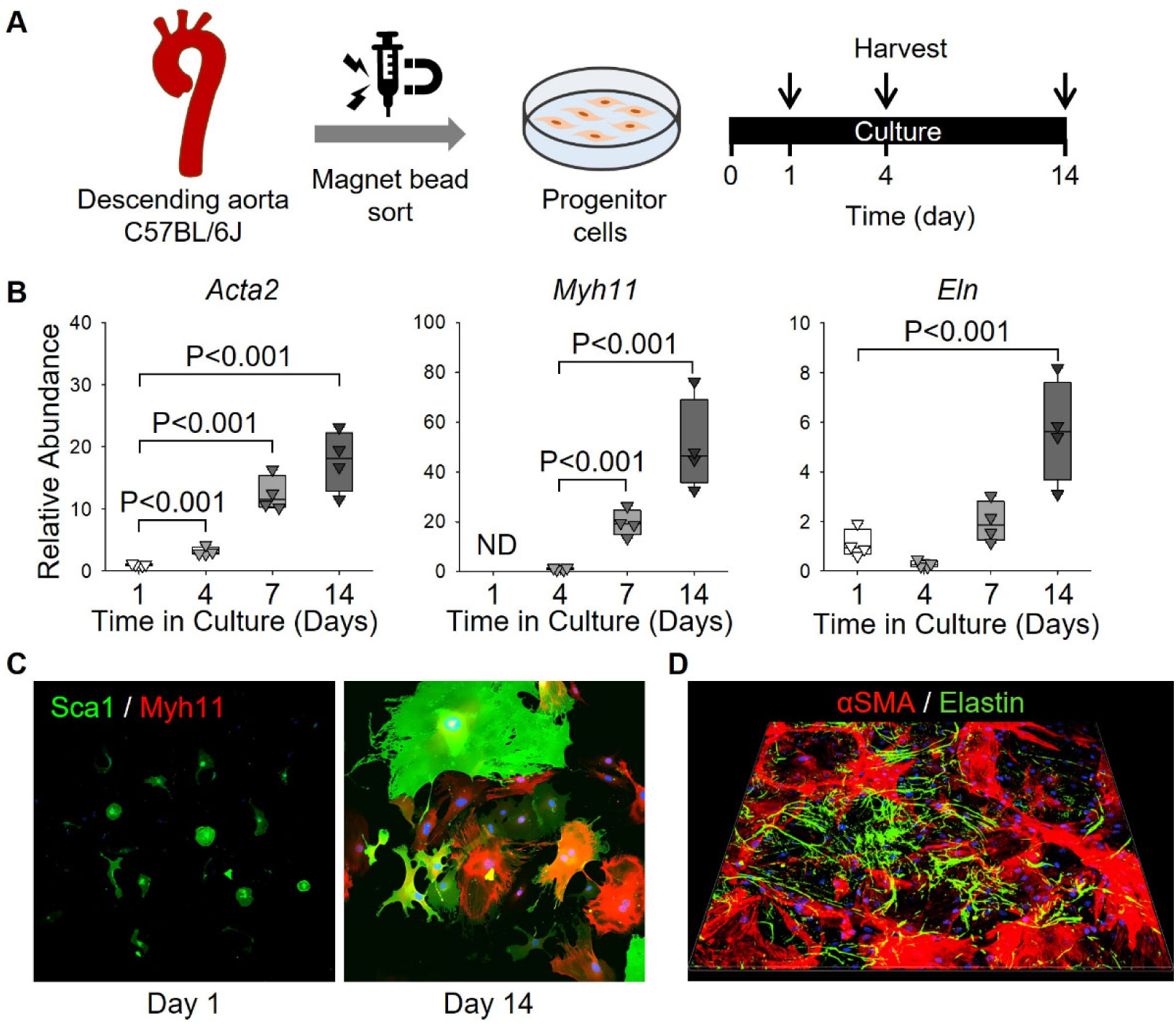
Acquisition of contractile and elastogenic phenotypes in adventitial progenitor cells. **(A)** Study design of in vitro experiments using adventitial progenitor cells. **(B)** Time-course qPCR analysis for *Acta2*, *Myh11*, and *Eln* mRNA abundance at day 1, 4, 7, and 14 in culture. N=4/group. ND indicates not detected. P values were determined by one-way ANOVA followed by the Holm-Sidak post hoc test. **(C)** Representative images of immunofluorescence for Sca1 (green) and MYH11 (red) in sorted Sca1+ cells at Day 1 and 14. **(D)** 3D reconstruction of immunostaining for αSMA and elastin.

## DISCUSSION

Our study provides several novel findings: (1) elastic fibers are regenerated in the false lumen wall of the descending aorta following AD; (2) although the newly formed elastic fibers differed structurally from those in the native medial layer, their presence associated with protection against rupture; and (3) de novo elastic fibers were produced by SMC-like cells that were not derived from resident medial SMCs, but originated from adventitial progenitor cells. Collectively, these findings offer new insights into the cellular and molecular basis of post-AD aortic reconstruction and may have important implications for the development of therapeutic strategies targeting the chronic sequelae of AD.

In the present study, a significant increase in *Eln* pre-mRNA was observed in chronic AD, in addition to increased *Eln* mRNA abundance. Although elastin expression in the aorta has been reported to be regulated at the post-transcriptional machinery (37–40), our findings suggest that transcriptional upregulation of elastin occurs in chronic AD. This raises the possibility that de novo elastin transcription following AD is driven by regulatory mechanisms distinct from those operating during normal developmental elastogenesis. We also demonstrated that adventitial progenitor cells acquired elastogenic ability after differentiation into SMC-like cells. Differentiation of vascular progenitor cells into SMCs has been reported to involve multiple signaling pathways, including TGF-β, Notch, and PDGF signaling (47, 58). To investigate the transcriptional regulation of elastic fiber-related genes and SMC-differentiation mechanisms, we performed spatial transcriptomic analysis. It revealed transcriptional differences in genes associated with TGF-β signaling between resident SMCs and elastogenic SMC-like cells. This finding raises the possibility that activation of this pathway contributes to the expression of elastic fiber genes and/or the differentiation of adventitial progenitor cells. In addition, given that epigenetic regulation may play a critical role in driving de novo elastin transcription and cell differentiation programs following AD, an integrative approach combining single-cell RNA sequencing and single-cell ATAC sequencing would be particularly powerful. This approach enables simultaneous profiling of transcriptional states and chromatin accessibility at single-cell resolution, thereby providing mechanistic insight into the epigenetic landscape underlying elastin synthesis and elastogenic cell differentiation in the false lumen wall.

Elastic fibers are essential structural components of the aortic wall, and their fragmentation is proposed to predispose to AD (22–24). In the present study, biomechanical analyses demonstrated increased wall stiffness in aortas with chronic AD, and de novo elastic fiber formation was absent in aortas that subsequently ruptured. These observations collectively suggest that elastic fiber regeneration may play a protective role against aortic rupture in the chronic phase after AD. Nevertheless, the mechanical properties of the aortic wall depend on multiple factors; in addition to elastic fibers, collagen fibers may also contribute (22, 59, 60). The biomechanical analyses performed in the present study reflect the composite stiffness of the whole vascular wall rather than the isolated contribution of de novo elastic fibers. Therefore, it remains unclear how and to what extent elastic and collagen fibers contribute to the observed mechanical changes. Future studies employing conditional genetic deletion of elastin or collagen fiber components will be needed to clarify their respective roles in the structural stability of the dissected aortic wall.

In this study, we used a *Gli1*-CreER^T2^ system for tracking adventitial progenitor cells. Gli1 is expressed in humans and is relatively specific for adventitial progenitor cell populations. In addition to Gli1, Sca1, and CD34 can be used as markers for adventitial progenitor cells. Sca1 is another widely used marker for mouse adventitial progenitor cells (47, 48, 56). However, there is no human ortholog, representing a fundamental limitation for translational studies (61). Also, Sca1 is observed in aortic endothelial cells. CD34 labels a broader population of adventitial progenitor cells but has limited specificity (61). In our studies using *Gli1*-CreER^T2^; *Rosa26*-mTmG mice, a small number of labeled cells were also detected in the aortic medial layer, raising the possibility of contamination by SMC-lineage cells. Therefore, no single marker perfectly defines adventitial progenitor cells, and interpretation based on a combination of markers and lineage-tracing approaches is necessary. A more fundamental challenge is that these Cre-based lineage tracing systems are designed for use in mouse models and cannot be directly applied to human AD tissue. Since lineage tracing is not feasible in human specimens, alternative strategies are required to infer cellular trajectories. Analysis of adventitial progenitor cell and SMC marker expression across multiple time points after AD, combined with trajectory-based pseudotime analyses using single-cell transcriptomic data, may provide a practical means of reconstructing the differentiation dynamics of adventitial progenitor cells in human tissues.

Despite advances in surgical and endovascular intervention, the long-term management of AD remains inadequate, and no pharmacological therapy has been established to inhibit vascular complications and directly promote false lumen stabilization beyond conventional blood pressure control. The present findings introduce several potential targets for therapeutic intervention. Therapeutic strategies aimed at directly augmenting de novo elastin synthesis could potentially enhance the mechanical integrity of the false lumen wall and reduce the risk of late rupture. In addition, promoting the expansion and differentiation of adventitial progenitor cells into elastogenic SMC-like cells represents a complementary therapeutic approach, as increasing the number of elastin-producing cells may further stabilize the dissected aortic wall.

In conclusion, the present study revealed that adventitial progenitor cells differentiate into SMC-like cells and produce de novo elastic fibers following AD. These findings establish a conceptual framework for targeting adventitial progenitor cells and elastin synthesis as novel therapeutic strategies to improve long-term outcomes in patients with AD.

## Supporting information

Supplemental Excel File for Data

## AUTHOR CONTRIBUTIONS

SI, PP, RW, and HS analyzed the data. SI and HS wrote the manuscript and prepared figures. SI, RW, and HS implemented experiments. TI and KO provided human samples. SI and HS contributed to the study design. All authors reviewed the manuscript.

## ACKNOWLEDGEMENTS

We thank Dr. Jay D. Humphrey and members of the Humphrey Laboratory at Yale University for their assistance with biomechanical testing and interpretation of the biomechanical data.

## SUPPLEMENTAL FIGURES

**Supplemental Figure 1.**
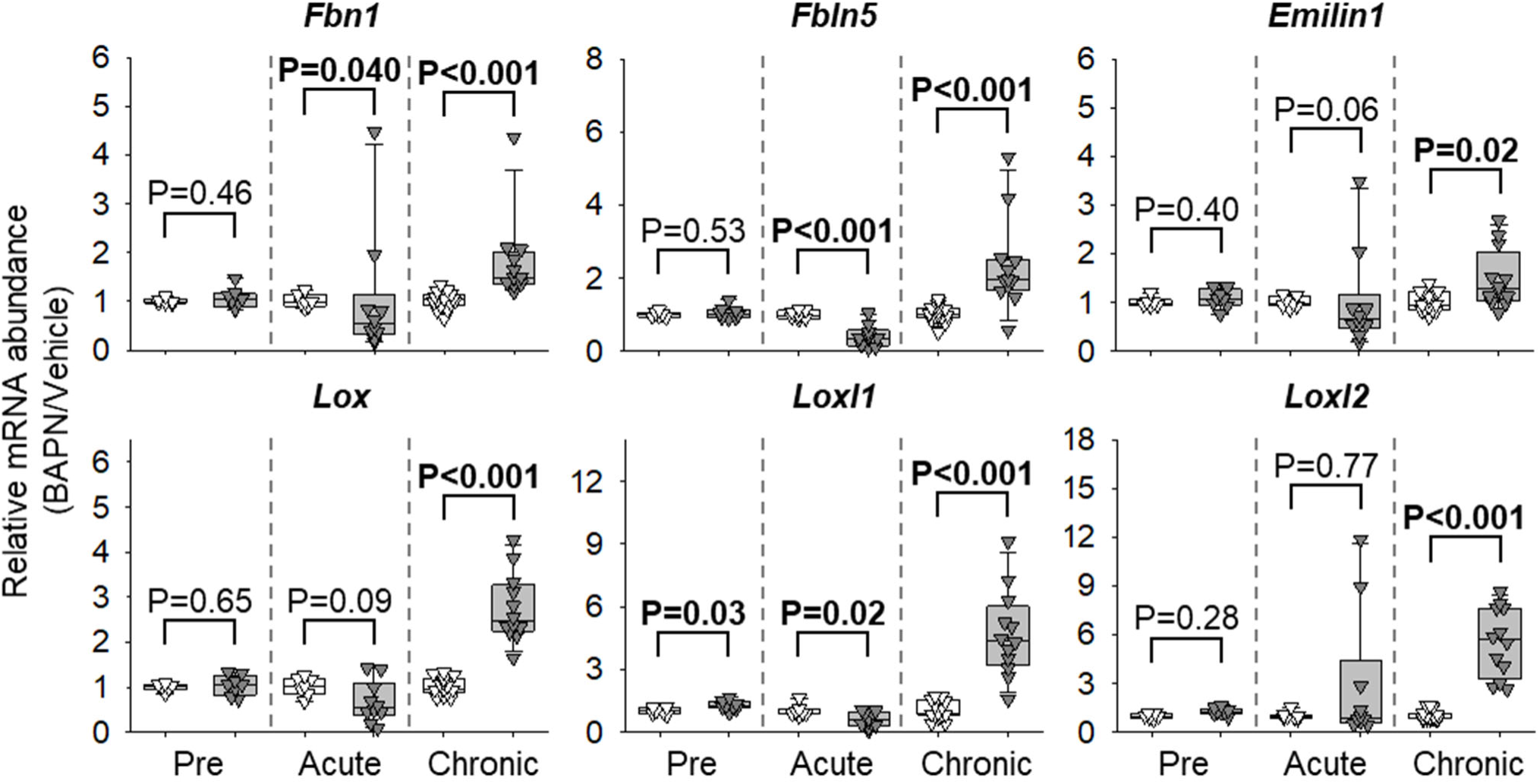
mRNA of elastic fiber-related genes increased in chronic AD of BAPN-administered mice. *Fbn1, Fbln5, Emilin1, Lox, Loxl1*, and *Loxl2* mRNA at pre, acute, and chronic phases after AD. N=6-15/group. P values were determined by Welch’s t-test or Mann-Whitney test between administrations at each time point.

**Supplemental Figure 2.**
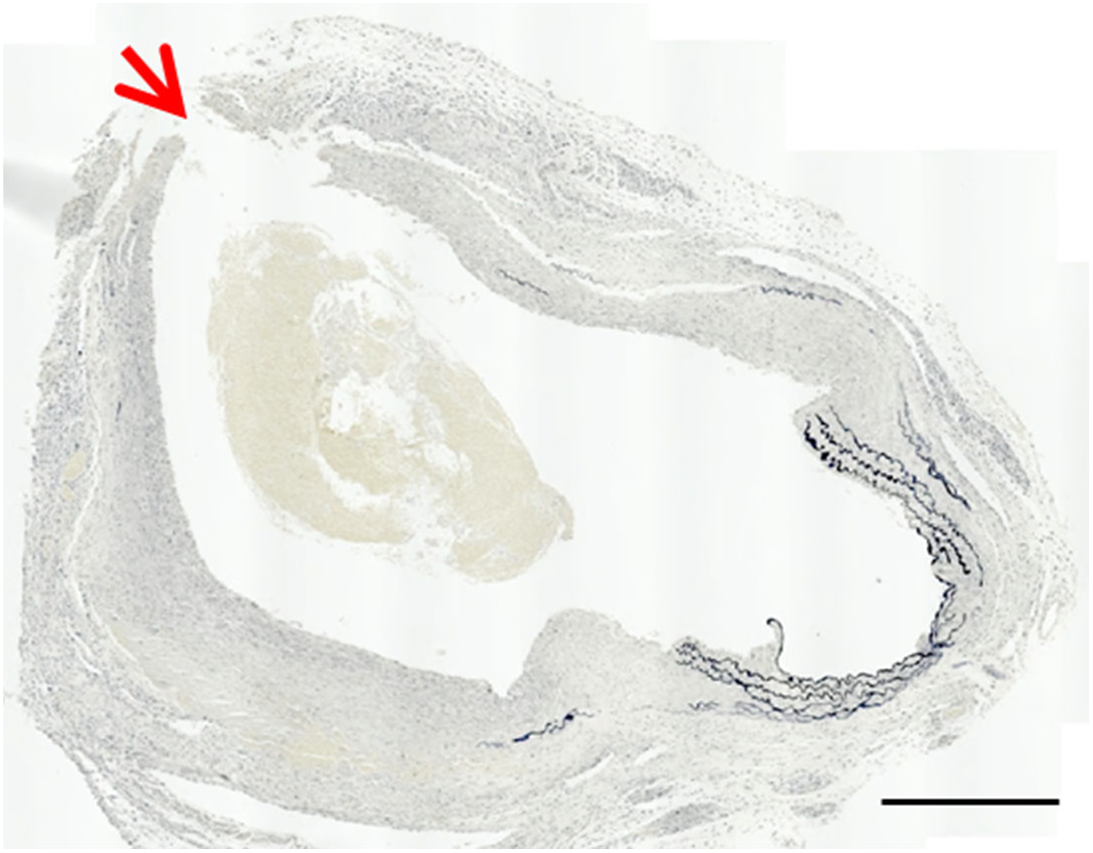
Aplasia of de novo elastic fibers in mice with late false lumen rupture. Representative image of Verhoeff iron hematoxylin staining in a mouse that died of false lumen rupture in the chronic phase following BAPN-induced AD. The red arrow indicates the rupture site. Scale bar = 500 µm.

**Supplemental Figure 3.**
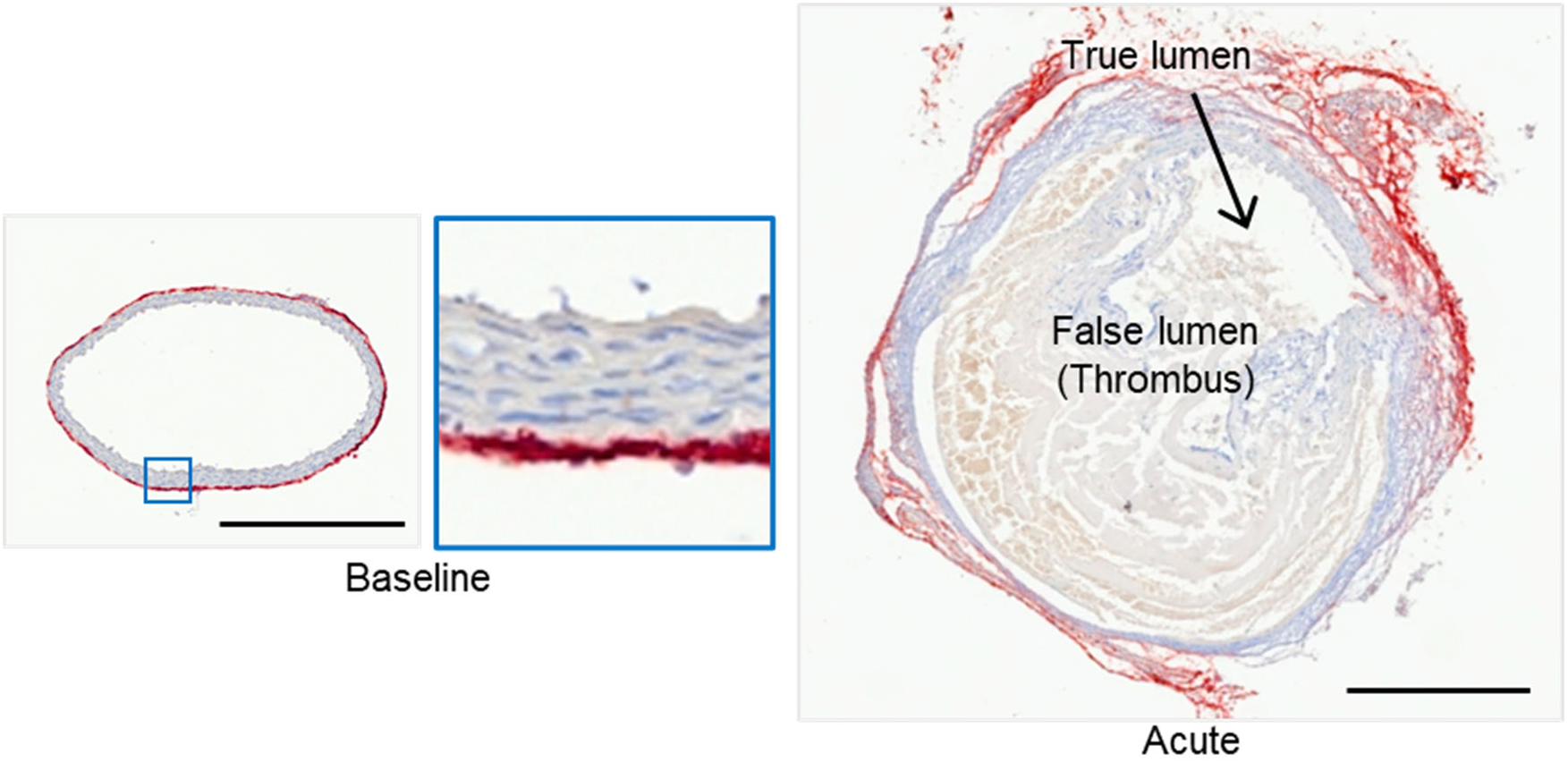
Distribution of Sca1+ vascular progenitor cells in the aortic wall at baseline and the acute phase following BAPN-induced AD in mice. Representative images of immunofluorescent staining for Sca1 in the descending aorta at baseline and the acute phase after BAPN-induced AD in mice. Scale bar = 250 µm.

**Supplemental Figure 4.**
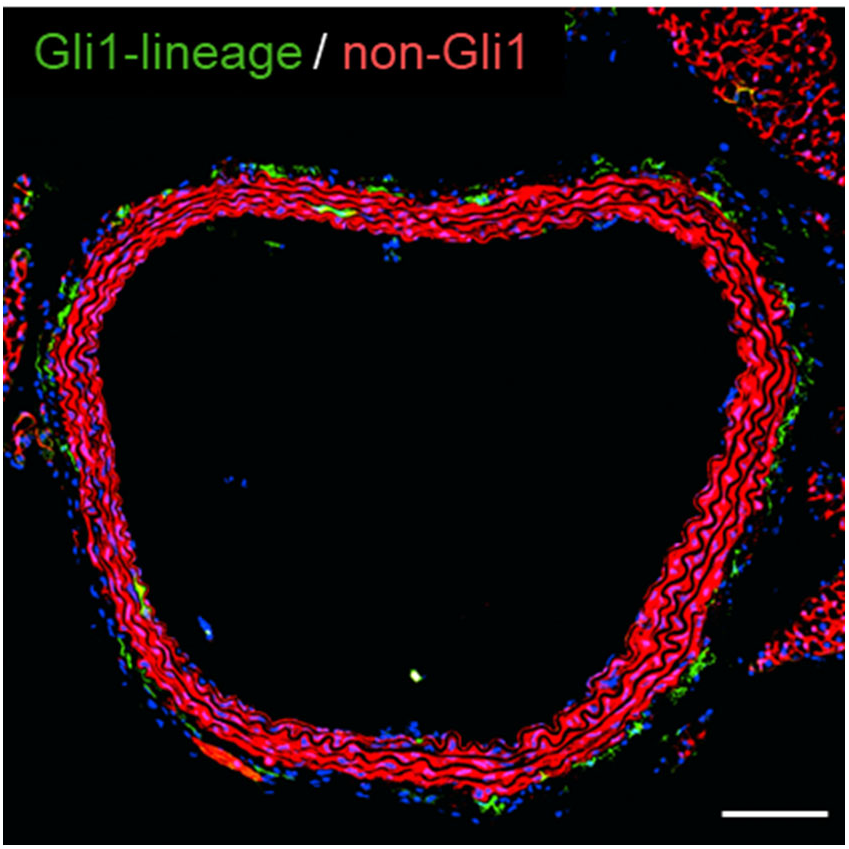
Gli1-lineage cells were detected mainly in the adventitia of the descending aorta from control mice. Representative aortic image of *Gli1*-CreER^T2^; *Rosa26*-mTmG mouse. Scale bar = 100 µm.

## MAJOR RESOURCE TABLE

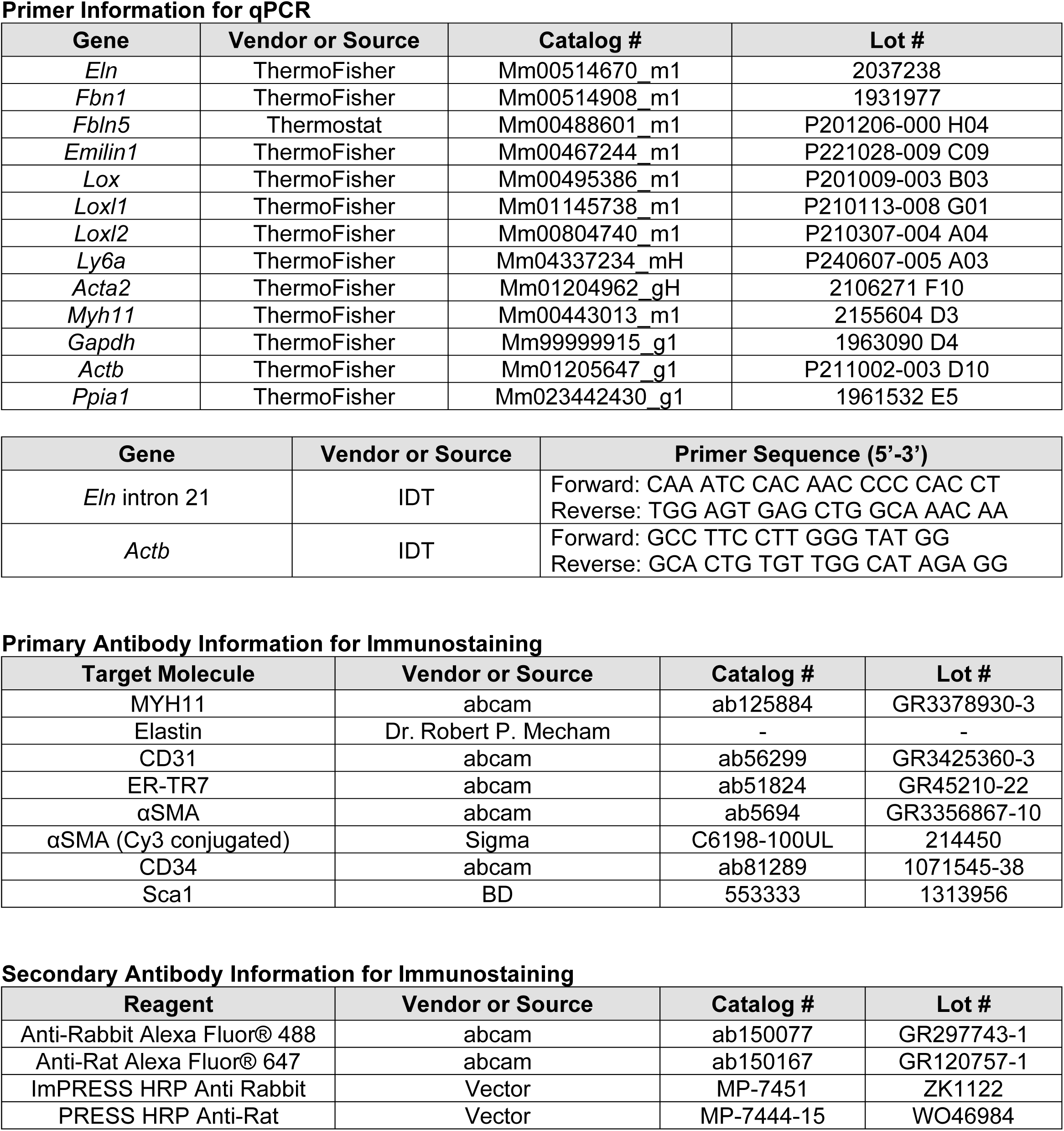

